# Fluctuating hydrodynamic forces increase fish locomotor variability in turbulence

**DOI:** 10.64898/2026.10.13.758844

**Authors:** Michael A. Calicchia, Divya Ramesh, Yangfan Zhang, Caleb Pipes, George V. Lauder, Rui Ni

## Abstract

Turbulence has been shown to alter fish locomotion and increase the energetic cost of swimming, yet the mechanisms underlying these effects remain unclear. To address this, we studied giant danios (*Devario aequipinnatus*) swimming at multiple speeds in both laminar and turbulent flows. Within this turbulent regime, mean tail-beat kinematics were largely unchanged relative to laminar conditions, but cycle-to-cycle variability increased, with larger effects occurring at higher speeds, where the velocity fluctuations are stronger. Turbulence also increased the frequency of asymmetric tail beats, particularly under stronger fluctuations, providing further evidence of increased locomotor variability. A quasi-steady hydrodynamic model suggests that velocity fluctuations in this flow regime produce relatively small changes in mean thrust and drag, whereas thrust variance scales with the square of velocity fluctuation magnitude. These variable forces likely prompt fish to adjust locomotor output to maintain position, which could explain the observed increase in kinematic variability. We further show that mechanical swimming power can be increased through correlations between fluctuating hydrodynamic forces and tail velocity. Together, these results suggest a mechanistic link between turbulent velocity fluctuations, locomotor variability, and increased energy expenditure.

## 1 Introduction

Aquatic environments are characterized by spatial and temporal variation in velocity and turbulence, creating a wide range of flow conditions through which animals such as fish must swim. Because these conditions affect locomotor responses, energy expenditure, and movement decisions, the physical mechanisms linking hydrodynamics to fish behavior have long attracted interest [1–9]. Extensive work has characterized the effects of mean flow, establishing that increased swimming speed requires changes in tail-beat frequency and amplitude [10–13] to generate the thrust needed to overcome fluid drag [14–17]. This is associated with elevated energy expenditure, which is expected to scale as *u*^3^, where *u* is the swimming speed [18–22].

Prior studies have also demonstrated that turbulence can further increase the fish’s energy expenditure [23–26]. However, it remains unclear how turbulence contributes to the elevated energetic cost [27]. Pavlov et al. proposed that turbulent fluctuations can increase the mean drag on the body, requiring greater kinematic effort to maintain position [28]. They suggested that the required power to overcome drag in turbulence should thus scale as *u*^3^ (1 + 3*I*^2^), where *I* is the turbulence intensity [28].

In a recent study, Zhang et al. [26] measured the total cost of transport of fish swimming in laminar and turbulent conditions over a variety of speeds. Given that the turbulence intensity in this study was approximately 20%, the drag-based scaling would predict a roughly 12% increase in the energetic cost, which is significantly less than the observed increase [26].

This suggests that additional mechanisms are at play. One possibility is that turbulence acts not only through changes in mean force, but also through fluctuations in hydrodynamic loading. Because turbulence is inherently stochastic, it can vary the thrust produced by the fish from one tail-beat cycle to the next by altering the relative velocity at the tail and the angle of attack between the caudal fin and the relative flow direction [29]. Such fluctuations may require cycle-to-cycle adjustments in locomotor output to maintain position and orientation, potentially increasing energetic cost even when mean force demand is unchanged.

To test whether this is a potential mechanism contributing to the increased cost of transport in turbulence, we first quantified how turbulence alters swimming kinematics. We recorded long-duration time series of fish swimming in laminar and turbulent flows over a range of swimming speeds. Our goal was to examine the kinematic effects of low to moderate turbulence (≈ 10% turbulence intensity or less) corresponding to moderate natural conditions commonly experienced by fishes rather than the extreme levels of turbulence present in many high-velocity streams. In this regime, mean hydrodynamic forces are not expected to increase substantially, suggesting that turbulence would act primarily through its fluctuating nature. The kinematic data were then used to estimate how the mechanical swimming power scales in turbulence, which should be directly proportional to the active metabolic rate [13, 29]. This approach provides a framework for linking turbulent velocity fluctuations to altered swimming kinematics and increased energetic cost.

## 2 Methods

### 2.1 Experimental setup

The experiments were performed in a recirculating water channel with a test section of 50 cm in length, 30 cm in width, and 30 cm in height. Temperature was maintained at 27.7 °C. Honeycomb baffles were installed at both ends of the test section to condition the inflow and prevent fish from leaving the measurement region during experiments. Fish were further confined to a smaller volume of the test section using a nearly-transparent cage made of fine net material to ensure they remained within the camera’s field of view for extended periods. This experimental arrangement was used previously by Zhang et al. [30] and by Ko et al. [31], who provide velocity calibrations that quantify the effect of the net walls on flow. By using porous walls, this experimental arrangement greatly reduced wall-effects and minimized the tendency of fish to swim near the cage wall.

Turbulence was generated using a passive grid made from Plexiglass (0.125 inch thickness) mounted upstream of the fish cage. The turbulence-generating grid consisted of an array of square openings (4 × 4), each measuring (1.5 × 1.5) cm, which corresponds to roughly 25% of the fish body length. To resolve the fish kinematics, a high-resolution streaming camera (AOS PROMON U1000) recorded ventral-view images of the fish at 200 fps. A front-surface mirror positioned at 45^°^ beneath the test section provided optical access to the ventral view. LED lights (Nila Zaila) were mounted above the experimental setup to illuminate the tank. A transparent acrylic plate was also placed on the water surface, covering the top surface area of the test section, to remove shadows from surface distortion. The entire setup was covered with laser blackout fabric (Nylon Fabric with Polyurethane Coating; Thorlabs Inc, New Jersey, USA) to minimize the effects of visual disturbances. A schematic of the setup is shown in Fig. 1(A).

**Figure 1.**
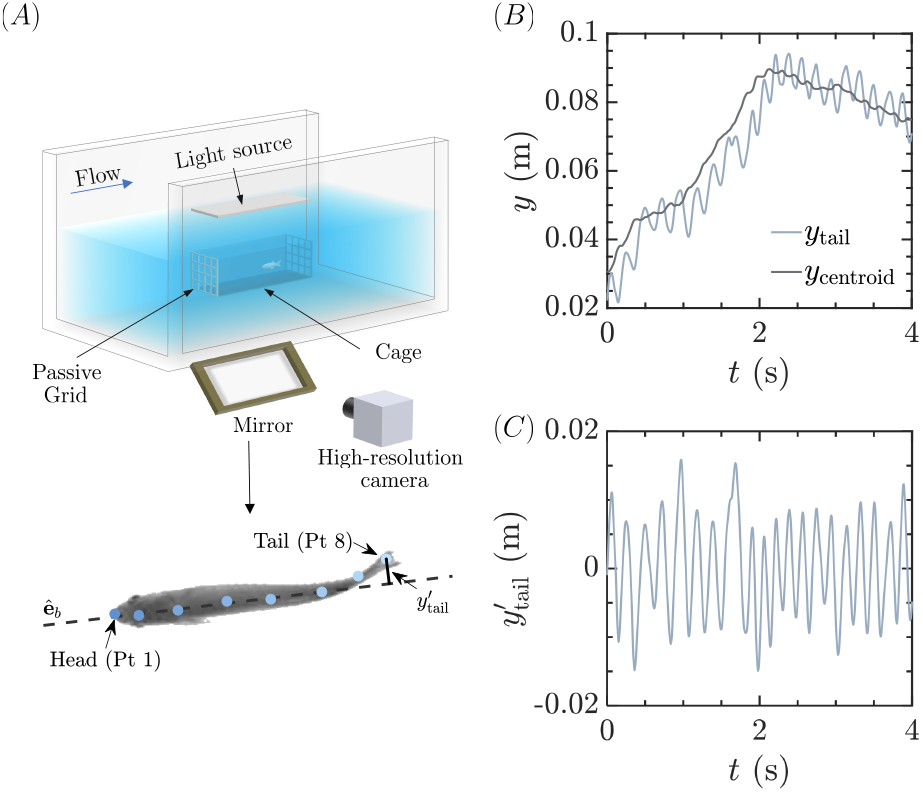
Experimental setup and quantification of fish motion. (A) Section of the flow tank highlighting major components including light source, high-resolution camera, fish cage, and the passive grid. An example of tracked fish midline points with the body-orientation unit vector **ê**_b_ and perpendicular tail displacement relative to the body axis, *y*^′^tail is also shown. (B) Example time series of the fish tail and centroid positions. (C) Example time series of the tail displacement perpendicular to the body orientation, independent of translational motion of the fish centroid.

### 2.2 Velocity measurements

This same streaming camera was used to record the motion of 50 *µ*m mean diameter plastic tracer particles at 1000 fps. The particles were illuminated within a single horizontal plane using a continuous-wave, diode-pumped, all-solid-state 532 nm green laser (5 W, MGL-N-532A, Opto Engine LLC). The recorded images were processed using Particle Image Velocimetry (PIV), where a multi-pass cross-correlation algorithm was applied to obtain the instantaneous velocity fields. The measurement region was centered on the cage mid-depth.

Velocity fields were acquired for both laminar and turbulent flow conditions using PIVlab V3.12 in MATLAB [32]. The reconstructed velocity fields were used to compute the mean flow velocity, turbulent intensity, and probability density functions of the streamwise and spanwise velocity components. The turbulence was further quantified by the characteristic length and time scales, as well as the energy dissipation rate *ε*.

To estimate these quantities, the commonly-used longitudinal (*D* _*LL*_ (*r*) = ⟨[*u*_*l*_ (*x*) − *u*_*l*_ (*x* + *r*)]^2^⟩ and transverse (*D* _*N N*_ (*r*) = ⟨[*u*_*n*_ (*x*) − *u*_*n*_ (*x* + *r*)]^2^⟩ structure functions were employed, where ⟨·⟩ denotes an average over all velocity pairs that have the same separation distance *r, x* is the spatial location, *u*_*l*_, *u*_*n*_ are the fluctuating velocity components projected into the longitudinal (along the direction of *r*) and transverse directions (perpendicular to the direction of *r*), respectively. The energy dissipation rate *ε* can be obtained from the structure functions using the following relationships (*D*_*LL*_ (*r*) = *C*_2_ (*εr*)^(2/3)^) and (*D* _*N N*_ (*r*) = 4/3*C*_2_ (*εr*)^(2/3)^), where *C*_2_ is a universal constant and is assumed to be 2 [33].

With this relationship, the structure functions can be used to obtain an approximation of the energy dissipation rate to estimate the integral 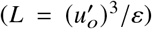 and Kolmogorov (*η* = *v*^3^ / *ε*^1/4^) length scales, where *v* is the kinematic viscosity of water and 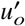 is the root-mean-square fluctuation velocity.

### 2.3 Experimental protocol

Giant danio (*Devario aequipinnatus*) was selected as the fish species for this study given the considerable previous research on locomotion for which this species has served as a model [26, 30, 31, 34–38]. Four individual fish were chosen randomly with body lengths ranging from 5.90 to 6.64 cm. Each fish was placed in the experimental setup the previous day, ~ 20 hours before the start of the trial, and made to swim against 1.0 BLs^−1^. The fish were allowed to get accustomed to 0.5 BLs^−1^ for the first 5 minutes before the start of the trial. Each fish swam in both laminar and turbulent conditions, with swimming speed incremented every 15 minutes from 0.5 to 6 BLs^−1^. The total duration of each trial was thus 1.25 hours. For this study, only speeds from 1 BLs^−1^ were used. Animal holding and experimental procedures were approved by the Harvard Animal Care IACUC Committee (protocol number 20-03-3).

### 2.4 Kinematic data processing

To resolve the fish’s kinematics, eight points along the midline, including the snout, peduncle, and tail, were tracked using DeepLabCut [39–41]. The marked points are shown in Fig. 1(A), with point 1 starting at the snout and point 8 ending at the tail. Due to the computational cost of the analysis, a one-minute subset of each fifteen-minute recording was used for tracking (note that we omitted the first five minutes because the fish was getting accustomed to the speed increase). The same selection criterion was applied consistently across trials to identify the interval in which the fish remained within the central region of the domain for the longest duration, minimizing boundary interactions and tracking loss. The tracked videos were then manually inspected to ensure accurate tracking of the midline points.

The primary kinematic quantity of interest was the displacement of the tail relative to the body midline, as tail motion is a primary contributor to thrust generation. Because the fish did not remain perfectly aligned with the swimming direction and frequently changed its lateral position within the test section, the tail location was quantified relative to the instantaneous body orientation rather than the fixed laboratory coordinate system. The third, fourth, and fifth midline points shown in Fig. 1(A) were used to determine the orientation of the fish relative to the swimming direction and define the instantaneous body orientation unit vector **ê**_*b*_. The relative tail position was then calculated by subtracting the center-of-mass position of the fish from the tail position. This removes the contribution of translational fish motion from the measured tail displacement.

Finally, the relative tail-position vector was projected onto the direction normal to the body orientation vector and is defined as 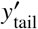. This provides a time series of the perpendicular displacement of the tail relative to the body midline, which was used to characterize the tail-beat kinematics. Examples of the tail-motion time series before and after removal of translational motion and perpendicular projection are shown in Fig. 1(B)-(C). Since the body length of each swimmer was measured and the instantaneous arc length from snout to tail can be calculated from the tracked midline points, the spatial calibration can be performed directly from the fish videos.

## 3 Results

### 3.1 Turbulent velocity fields

The p.d.f.s of the streamwise *u* and spanwise *v* velocities are shown in Fig. 2(A)–(B). The shifting peak in the streamwise velocity p.d.f. demonstrates that the mean flow velocity increases. Furthermore, the widening of both the streamwise and spanwise velocity distributions with increasing mean flow indicates that the velocity fluctuations increase proportionally. This behavior is expected because the turbulence is coupled to the mean flow velocity.

**Figure 2.**
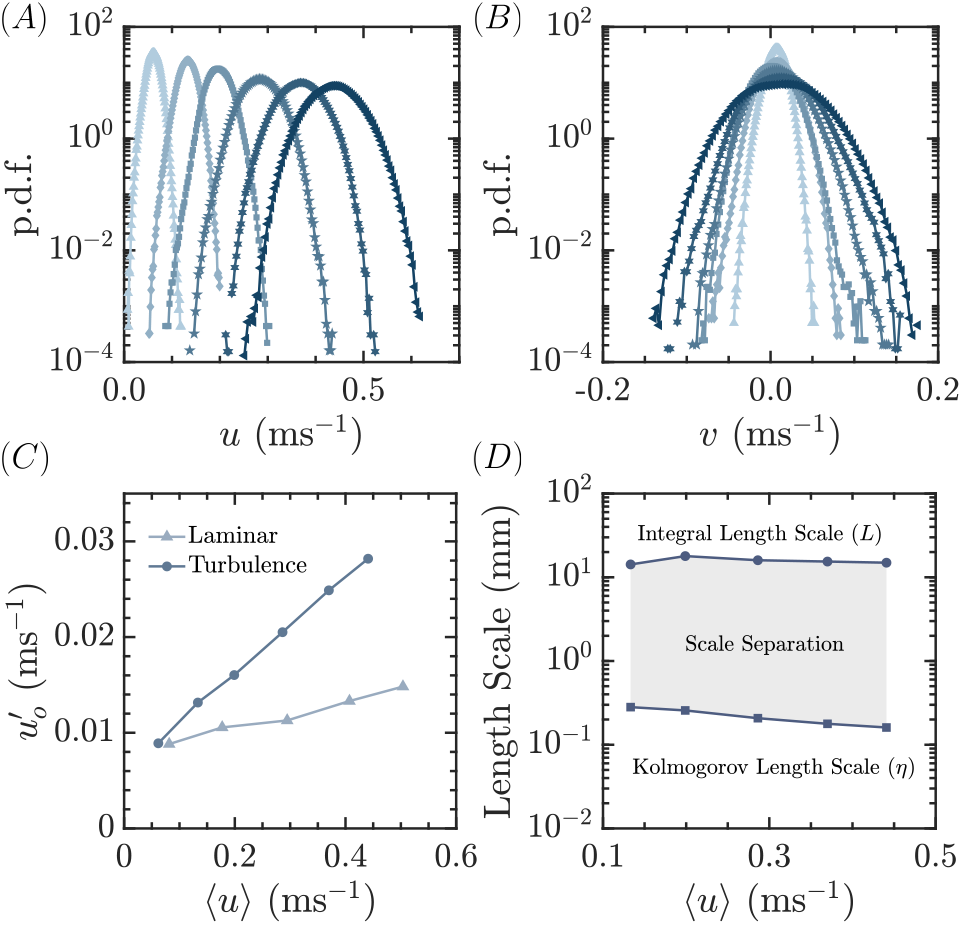
Characterization of the turbulence generated by the passive grid. Probability density functions of the (A) streamwise *u* and (B) spanwise *v* velocity for each swimming speed. (C) Root-mean-square velocity as a function of the swimming speed for laminar and turbulent conditions. (D) Estimates of the scale separation present in the turbulent flow demonstrating how the integral scale is roughly constant while the Kolmogorov scale decreases with increasing flow speed.

Fig. 2(C) shows the root-mean-square (RMS) velocity fluctuation magnitude 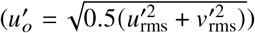 as a function of swimming speed, for the laminar and turbulent cases. Here 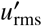 is the RMS of the streamwise velocity fluctuations, and 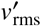 is the RMS of the spanwise velocity fluctuations. As expected, the passive grid increased the magnitude of velocity fluctuations and the rate of increase of fluctuations with swimming speed. The turbulent cases exhibit a turbulence intensity 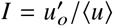 of approximately 7%, whereas the laminar flow conditions have a turbulence intensity of approximately 3%, which is inherent to the setup.

Using the two-dimensional structure function analysis described previously, the energy dissipation rate was estimated and used to determine the integral and Kolmogorov length scales. Fig. 2(D) shows the scale separation present within the turbulent flow fields. The integral length scale remains approximately constant (*L* ≈ 1.5 cm), consistent with the flow fields generated in Zhang et al. [26]. However, because the turbulence generated in this setup was weaker, the scale separation is reduced due to the larger observed Kolmogorov length scale.

Nevertheless, the flow remains within the same scaling regime where the turbulent length and momentum scales are small compared with those of the fish across all speeds. Specifically, the ratios of the integral length scale to fish body length and turbulent to fish momentum flux remain well below the thresholds identified in literature suggesting that the turbulence should not significantly affect a fish’s ability to maintain postural control [5, 42–44].

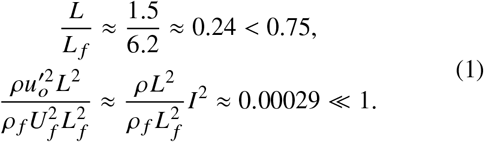

Here, the momentum flux of the largest eddies is expressed as 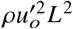, and that of the fish as 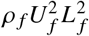, where *ρ* is the density of water, 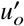 is the RMS fluctuation velocity, *L* is the integral length scale, *ρ* _*f*_ is the density of the fish, *U* _*f*_ is the swimming speed, and *L* _*f*_ is the body length.

### 3.2 Kinematic variations in turbulence

Next, we interpret the kinematic results in the context of the turbulence-induced hydrodynamic forces. The increase in the mean drag force due to turbulent velocity fluctuations can be estimated by decomposing the local velocity into mean and fluctuating components, *u* = ⟨*u*⟩ + *u*^′^, and expanding the quadratic drag scaling

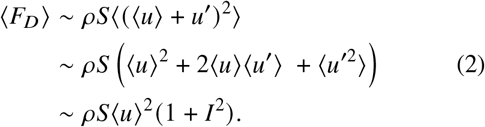

Here *ρ* is the fluid density, and *S* is the surface area. Given that the turbulence intensity was 7%, the increase in the drag force is estimated to be approximately 0.5%.

As mentioned earlier, turbulence may also influence the mean thrust production by modifying the relative velocity experienced at the tail and the angle of attack between the caudal fin and the relative flow direction. Since the majority of thrust production for this species originates from the caudal fin and is primarily associated with the leading-edge vortex (LEV) [45], a quasi-steady lift-based model can be used to estimate the scaling of thrust production. Following the approach of Seo et al. [29], the thrust force is given by

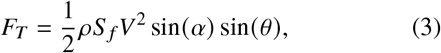

where *S* _*f*_ is the caudal fin area, and *V* is the relative velocity at the tail defined as, 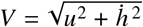. Here, *u* is the swimming speed, and 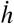 is the tail heaving velocity, which scales with the product of the tail-beat amplitude and frequency (*A f*). The instantaneous effective angle of attack is defined as 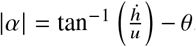, where *θ* is the pitching angle of the tail.

In turbulent flows, both the relative velocity and effective angle of attack contain contributions from the velocity fluctuations. These quantities can therefore be expressed as the sum of mean and fluctuating component. Here, we decompose each relevant variable into a phase-locked, cycle-averaged mean 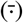 and a fluctuating component. The instantaneous thrust force is thus expressed as a function of phase *ϕ* and cycle *k*:

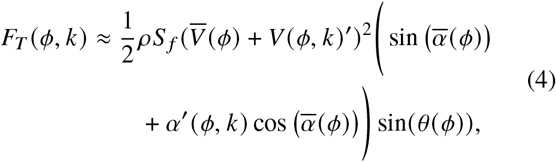

where a first-order Taylor expansion is used to linearize the sin 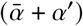 term. By expanding the relative velocity term, taking the cycle average, neglecting higher-order and covariance terms, and then phase-averaging (⟨·⟩_*ϕ*_), the scaling for the total average thrust force considering only the influence of turbulence can be expressed as:

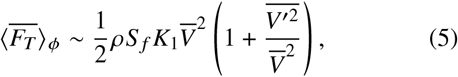

where *K*_1_ is an order one constant resulting from the phase-averaging.

The fluctuating relative velocity term 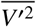 can be approximated as 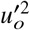 using a first-order expansion of the relative velocity and assuming comparable contributions from the turbulent velocity components. Therefore, assuming 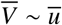 the effect of turbulence on mean thrust production scales with the square of the turbulence intensity, similar to the scaling of mean drag. Further details of the derivation of the mean thrust scaling are provided in Appendix A.

This analysis suggests that turbulence-induced changes in the mean drag and thrust forces are relatively small and of comparable magnitudes. Therefore, fish should not need to significantly modify their mean tail-beat kinematics to compensate for the additional hydrodynamic forces imposed by turbulence. Consequently, the mean tail-beat kinematics, characterized by *A f*, are expected to remain approximately unchanged between laminar and turbulent conditions.

To test this expectation, Fig. 3 shows the probability density functions (p.d.f.s) of *A f* normalized by the fish body length *L* _*f*_ for the four swimming speeds tested. The results are aggregated across all swimmers. The smoothness of the p.d.f.s indicates that a sufficient number of tail beats were captured in the 1 min of analyzed data.

**Figure 3.**
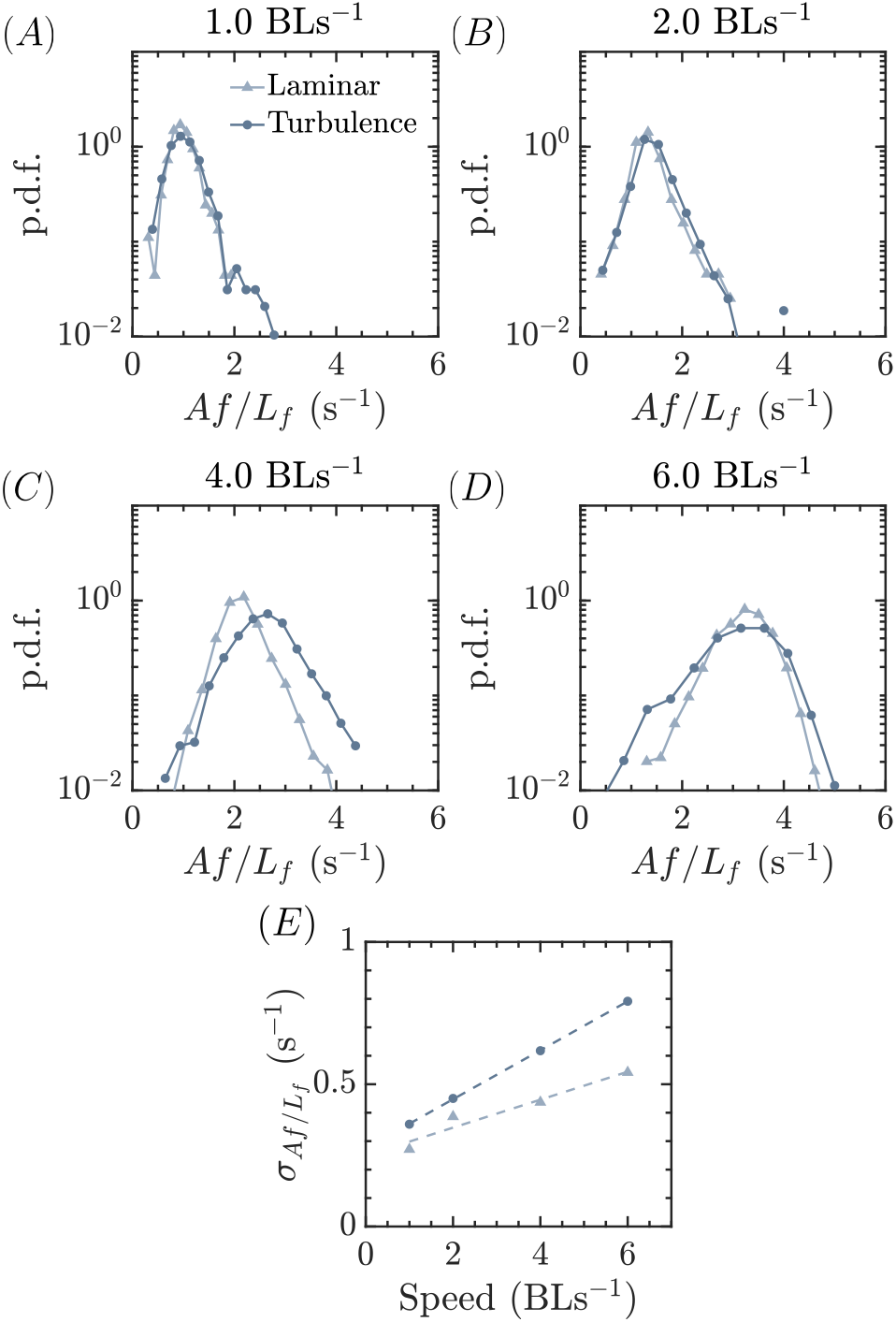
Variability of tail-beat kinematics in laminar and turbulent flows. Probability density functions of the product of tail-beat amplitude and frequency normalized by fish body length, for laminar and turbulent conditions at (A) 1.0 BLs^−1^, (B) 2.0 BLs^−1^, (C) 4.0 BLs^−1^, and (D) 6.0 BLs^−1^. (E) Standard deviation of tail-beat kinematics as a function of swimming speed and turbulence level, with dashed lines representing linear fits to the data points.

For the majority of swimming speeds, the peak of the *A f* / *L* _*f*_ distribution remains relatively unchanged between laminar and turbulent conditions, indicating similar mean tail-beat kinematics. The largest deviation in the peak value occurs at 4.0 BLs^−1^. However, as swimming speed increases and the magnitude of the turbulent velocity fluctuations rises, the distributions of *A f* / *L* _*f*_ become broader, suggesting greater cycle-to-cycle variability in tail-beat kinematics.

To better understand this observation, we next consider fluctuations in thrust production resulting from the surrounding turbulent flow. Utilizing Eq. 3, a first-order Taylor expansion can be used to express deviations from the mean thrust force due to turbulent velocity fluctuations as a function of both the phase and cycle:

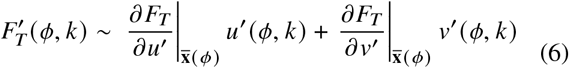

Here, the derivatives are evaluated at the mean state 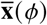. By squaring this expression, taking the cycle-average, assuming comparable contributions from the turbulent velocity components, and then taking the phase-average, the variance in thrust from cycle to cycle can be approximated as

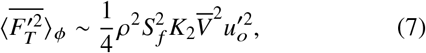

where *K*_2_ is another order one constant resulting from the phase-averaging. A more detailed derivation of this expression is provided in Appendix A.

This result indicates that the thrust variance scales with the square of the turbulent velocity fluctuations. Therefore, as swimming speed increases, larger velocity fluctuations are expected to generate larger variation in thrust production, which may cause fish to adjust locomotor output to maintain position and orientation.

The results in Fig. 3(E), which plots the standard deviation of *A f* / *L* _*f*_ as a function of swimming speed, are consistent with this interpretation. Tail-beat variability increases with swimming speed under both laminar and turbulent conditions. However, tail-beat kinematics under turbulent conditions consistently exhibit greater variability, with the standard deviation increasing more rapidly with swimming speed than in laminar conditions. This is likely due to the larger contribution of the 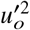 term in the thrust variance scaling under turbulence.

To further investigate the increase in kinematic variability observed in turbulence, a stroke asymmetry index *S* was defined for each tail beat *k* to quantify how turbulence influences the left-right symmetry of tail-beat kinematics.

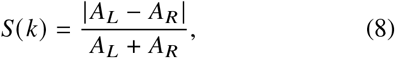

where cycles were defined from minimum-to-minimum tail position. The left-side amplitude, *A*_*L*_, was calculated as the difference between the cycle maximum and the left-side minimum, while the right-side amplitude, *A*_*R*_, was calculated as the difference between the cycle maximum and the right-side minimum. This approach avoids potential errors associated with zero-crossing detection resulting from variations in body orientation and the resulting projection of the perpendicular tail position.

The probability density functions of the stroke asymmetry index for all flow conditions are shown in Fig. 4. At lower swimming speeds and weaker velocity fluctuations, the distributions are characterized by a high probability of symmetric tail beats, with long distribution tails indicating occasionally larger asymmetric tail motions. The distributions are similar between laminar and turbulent conditions, suggesting that turbulence has a limited influence on tail-beat asymmetry at lower speeds.

**Figure 4.**
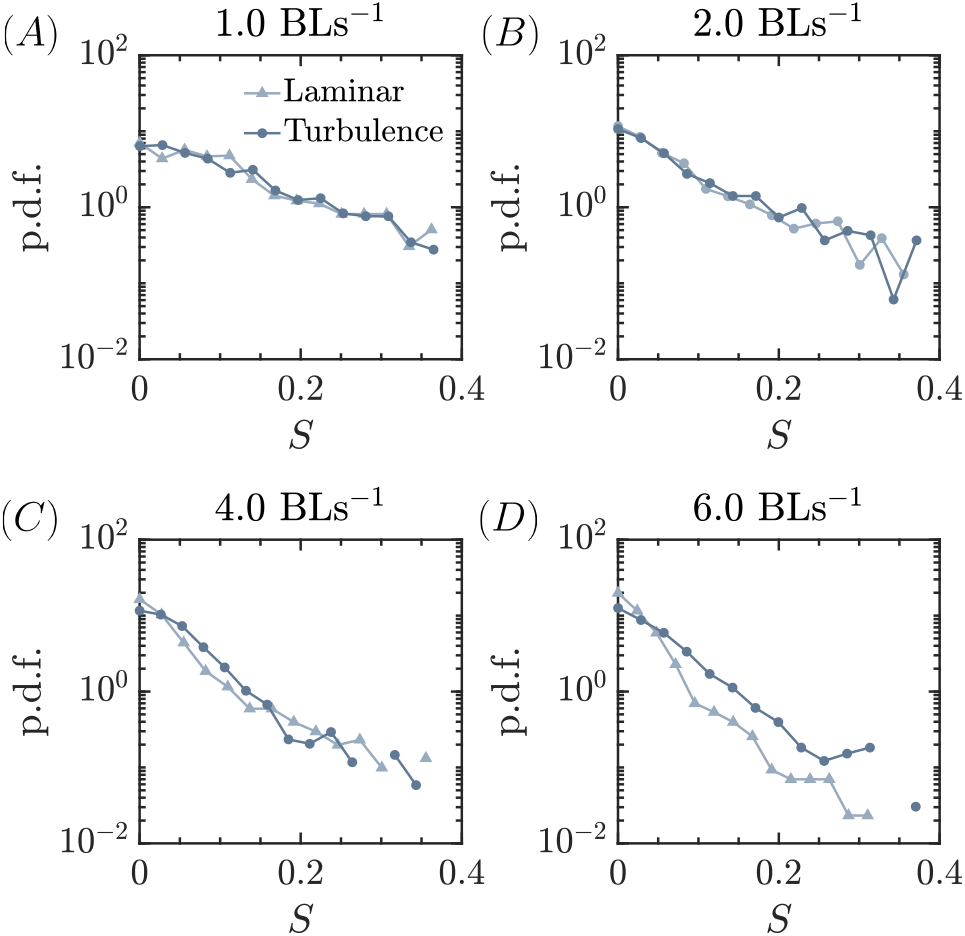
Variation in tail-beat asymmetry as a function of swimming speed and turbulence. Probability density functions of the stroke asymmetry index in laminar and turbulent conditions at (A) 1.0 BLs^−1^, (B) 2.0 BLs^−1^, (C) 4.0 BLs^−1^, and (D) 6.0 BLs^−1^.

At higher swimming speeds and larger velocity fluctuations, the distributions continue to exhibit a preference for symmetric tail beats but show increased differences between laminar and turbulent conditions. In particular, turbulence increases the occurrence of small-to-moderate asymmetric tail beats. This increase in asymmetry may result from active adjustments in thrust production as fish respond to local flow variations or from corrective turning moments used to maintain position and orientation [37, 46].

Regardless of the underlying mechanism, these results show that turbulence is associated with increased asymmetry in tail-beat kinematics, providing further evidence that stronger flow fluctuations produce greater variability in locomotor output.

### 3.3 Implications for the energetic cost of locomotion

Finally, we examine how variations in force and kinematics may influence the energetic cost of swimming. Following Seo et al. [29], the power required to move the tail laterally can be approximated as 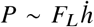, where the lateral force can be expressed as

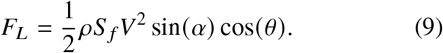

Because turbulence influences both the hydrodynamic forces and the fish’s kinematics, the lateral force and tail velocity are decomposed into the sum of a phase-locked, cycle-averaged mean and a fluctuating component. Substituting these decompositions into the power expression yields

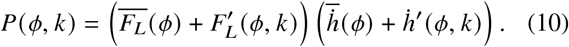

Next, we obtain the obtain the mean power at each phase by averaging over the cycles. Assuming the averaged fluctuations vanish, the expression for the average power at each phase is given as:

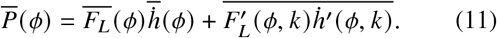

This expression shows that fluctuations in lateral force and tail velocity contribute to the mean power consumption at each phase through their covariance. Thus, even if the mean force and mean kinematics remain approximately unchanged, correlated fluctuations between force and tail velocity can increase the mean power consumption.

Lastly, to obtain the total power, the expression is then phase averaged.

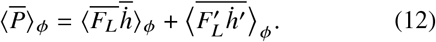

To determine whether the fluctuating term is expected to contribute positively to power consumption, we first consider the sources of lateral force variations: fluctuations due to turbulence, modulations of the tail pitching angle, and variations in the tail velocity. A first-order Taylor expansion is then used to approximate the fluctuations in the lateral force from these sources. Substituting this expansion into the covariance term in Eq. 11 yields

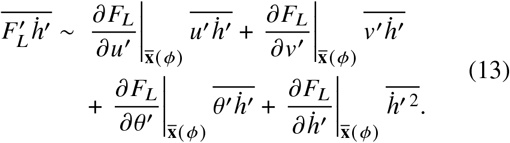

Here, (*ϕ, k*) has been dropped for simplicity.

It is difficult to predict both the magnitude and sign of the covariance terms in Eq. 13 without knowing exactly how the fish responds to the surrounding flow. However, perfect correlation between turbulent velocity fluctuations and the fish’s response would require the fish to sense and respond to changes in the incoming flow without a phase delay, which may not always be possible. Moreover, fish do not respond to turbulent fluctuations as purely efficiency-driven systems. Their responses reflect multiple behavioral demands, and therefore it is equally plausible that in certain situations the fish’s response may be negatively correlated with, or even independent of, the surrounding flow fluctuations. Thus, it is likely that first two covariance terms will be small on average.

The third term represents the covariance between tail pitch angle and tail velocity. Both *θ*^′^ and 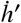 are actively controlled by the fish, which may adjust these quantities in different combinations in response to turbulent fluctuations. As a result, the sign and magnitude of 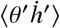 cannot be readily predicted, and thus this term is ignored for simplicity. In contrast, in the final term, 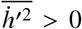, always. Thus, if the derivative is always likely to be positive, then, cycle-to-cycle variability in tail velocity will always increase the mean mechanical power required to generate lateral force and ultimately thrust.

By assuming that the covariance terms are negligible, substituting the derivative into Eq. 13, and using the relationship that 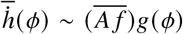, the covariance contribution to the average power at each phase can be obtained:

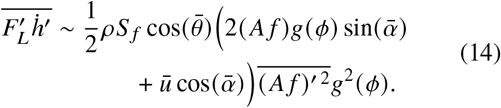

Here, *g* (*ϕ*) is an oscillatory function bounded by [−1, 1]. A more detailed derivation of this term and further discussion as to why it is expected to be positive is provided in Appendix B.

By averaging this expression over the phase as well as the mean component in Eq. 12, the scaling for the total average power can be expressed as:

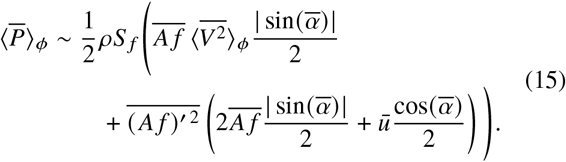

Fig. 5(A) demonstrates the expected scaling of the covariance term in Eq. 14 when the parameters of the present study are applied. The covariance term increases more rapidly with swimming speed in turbulence than in laminar flow, due to the coupling between larger turbulent velocity fluctuations and increased kinematic variability.

**Figure 5.**
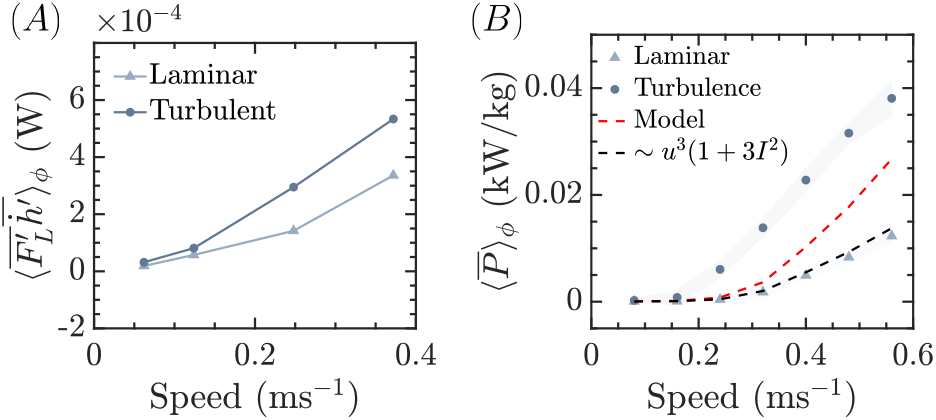
Fluctuations in fish kinematics and hydrodynamic forces lead to increased energy expenditure. (A) Scaling of the covariance term in the power equation as a function of swimming speed for laminar and turbulent conditions. (B) Metabolic power as a function of swimming speed for laminar and turbulent conditions, extracted from Zhang et al. [26], with the model prediction overlaid in red.

Next, we look to extend this scaling to the metabolic measurements presented in Zhang et al. [26] by making the following assumptions. First, it is assumed that the metabolic power will scale with the mechanical swimming power. Second, it is assumed that the variability in *A f* scales similarly with the magnitude of the turbulent velocity fluctuations 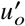, and the mean *A f* scales similarly with swimming speed.

The model is first used to predict the relative increase in power associated with turbulence compared with laminar flow. This predicted increase is then applied to the laminar metabolic power curve to estimate the turbulent metabolic power. The resulting prediction is shown by the dashed red line in Fig. 5(B). Because the model is a first-order, quasi-steady approximation, assumes all covariance terms are negligible, and neglects additional energetic costs such as added-mass effects and the muscular cost of moving the body in turbulence, exact agreement with the measured turbulent metabolic power is neither expected nor obtained.

Nevertheless, the model reproduces the observed trend that the energetic penalty associated with turbulence increases with swimming speed and turbulent fluctuation magnitude. The agreement between the trend of the model and experimental results suggests that the coupling between turbulent force fluctuations and cycle-to-cycle variability in tail-beat kinematics represents a plausible mechanism contributing to the increased energetic cost of swimming in turbulence. Importantly, this model scales power based on the fish’s response to turbulence, rather than turbulence intensity alone, and therefore captures the experimental trend more accurately than the previously proposed drag-based model (dashed-line prediction).

## 4 Conclusions

In this study, we found that mean fish kinematics were largely unchanged by turbulence levels of about 7%, whereas tail-beat variability and asymmetry increased. These observations are consistent with the expected scaling of mean and fluctuating thrust forces. Because the turbulence intensity was relatively low, average thrust production was not expected to change substantially. However, larger velocity fluctuations likely generated greater thrust deviation, which in turn increased cycle-to-cycle variability in locomotor output. Our results further suggest that these fluctuating forces, together with the associated kinematic adjustments, may contribute to the elevated energetic cost of swimming in turbulence, particularly at higher swimming speeds.

It is important to note that, in these experiments, the characteristic length and momentum scales of the turbulence were smaller than those of the fish. As a result, the turbulence primarily introduced fluctuations in hydrodynamic forcing without markedly altering mean force requirements or mean swimming kinematics. In flow regimes where the turbulent scales are more comparable to those of the fish, however, the effects on locomotion may be substantially larger, potentially influencing both mean kinematics and cycle-to-cycle variability and, consequently, having a greater effect on the swimming energetics.

Thus, fish–turbulence interactions may depend on several non-dimensional parameters, including *L*/*L* _*f*_, 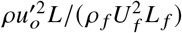, and 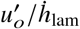, which help indicate whether postural instability is expected, whether mean hydrodynamic loading is altered, and how strongly fish must modulate kinematics to compensate for fluctuating thrust.

To more effectively study these regimes, future work should aim to control the turbulence independently of the mean flow velocity. This could be achieved using active turbulence-generation methods, such as a jet array, that inject energy into the flow rather than extracting it from the mean current, as is the case with passive grids [27]. Such experiments would provide a powerful test of how various turbulent environments can impact locomotor performance and energetic cost.

Furthermore, simultaneous measurements of fish–turbulence interactions would improve our understanding of how fish respond to turbulent flow and may enable better predictions of the covariance terms that were ignored in this study. These measurements could also make it possible to resolve surface pressure and ultimately hydrodynamic forces on the body [47, 48] and thus the forces exerted by the body, which may be more closely linked to energetic costs.

## A Appendix 1: Derivation of Thrust Scaling

In this section, the detailed derivation of the mean and fluctuating thrust scaling due to turbulence is provided. Eq. 4, shows the expression for the thrust in turbulence as a function of phase *ϕ* and cycle *k*. Expanding this expression and taking the phase-locked, cycle average yields the following assuming the averages of the fluctuating terms are zero:

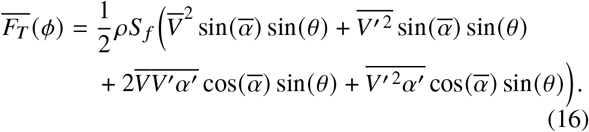

By ignoring the covariance term and the higher order fluctuating components, and phase-averaging, the total thrust per cycle can be expressed as

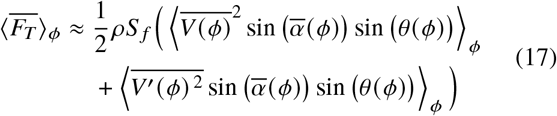

The magnitude of each term will be set by 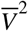 and 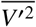 respectively and a first order positive pre-factor *K*_1_ as a result of the phase averaging. Thus, the average thrust over a cycle can be expressed as shown in Eq. 5.

To approximate the magnitude of the 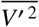, a first-order Taylor expansion of the relative velocity is used where 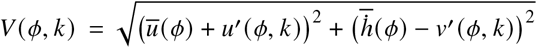, and the derivatives are evaluated at the mean values:

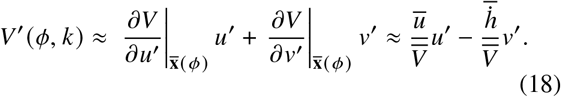

Next, the expression is squared, then cycle-averaged yielding:

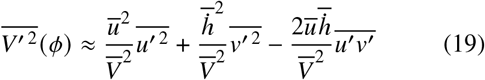

Given that the turbulence generated by the passive grid is nearly isotropic, it is reasonable to assume that the fluctuations in each direction are of comparable magnitude and that the covariance of the velocity fluctuations is small, which results in the following:

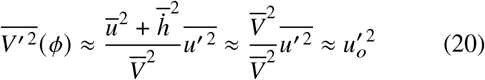

Next, to estimate the fluctuations of the thrust force due solely to the turbulence a first-order Taylor expansion is again utilized, as shown in Eq. 6. The variance of the thrust force at each phase is then obtained by squaring this expression, and taking the cycle average:

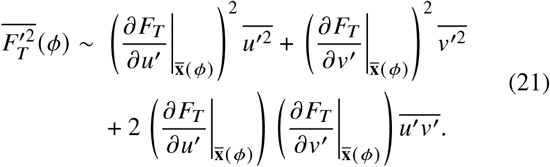

By assuming that the fluctuating components have similar magnitude and the covariance term is approximately zero, the equation reduces to

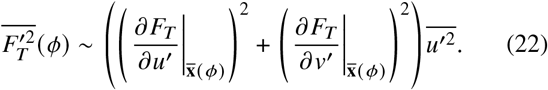

The derivatives in this expression are shown below:

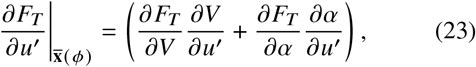

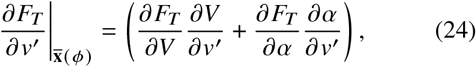

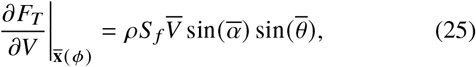

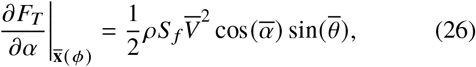

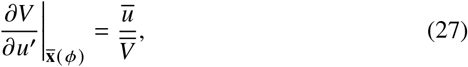

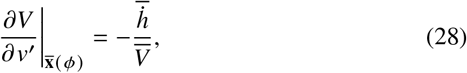

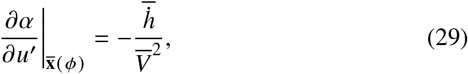

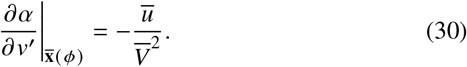

Substituting these derivatives into Eq. 22 yields:

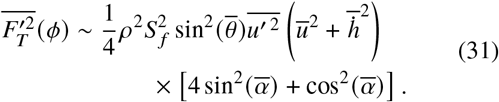

By noting that 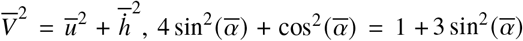, and averaging over the phase, the total thrust variance can be expressed as

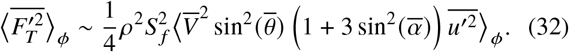

The phase average yields a scaling of the form 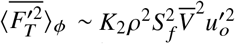, where *K*_2_ is an order-one coefficient, as shown in Eq. 7.

## B Appendix 2: Derivation of Power Scaling

In this section, the detailed derivation of the power scaling is provided. Utilizing a similar approach as before, a first-order Taylor expansion is used to approximate the fluctuations of the lateral force due to turbulence as well as the variations in the tail velocity and tail pitching angle. Under the assumption that the covariance terms in Eq. 13 are negligible, only the following derivatives are necessary to evaluate it:

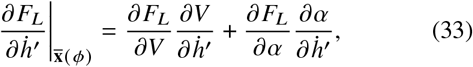

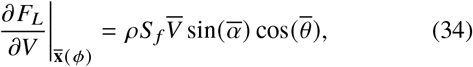

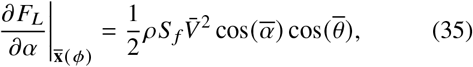

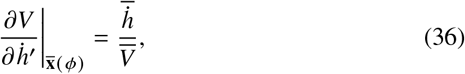

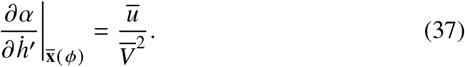

Substituting these expressions into the last term of Eq. 13 yields, where the average quantities are still a function of *ϕ*:

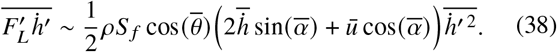

It is next assumed that 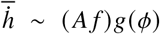 and thus 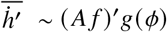, where *g* (*ϕ*) is an oscillatory function bounded from [−1, 1]. Substituting this into Eq. 38, then yields Eq. 14

Next, we consider whether this equation is expected to be positive at each phase. Given that 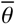 and 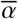 are expected to be bounded by [−*π*/2, *π*/2], the cosine terms in Eq. 14 are nonnegative. Furthermore, 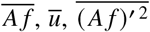 and *g*^2^ (*ϕ*) are always positive. The only remaining phase-dependent sign contribution is 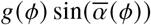. During rightward tail motion, *g* (*ϕ*) is positive and 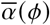 is also predominantly positive. Likewise, during leftward motion, *g* (*ϕ*) is negative and 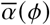 is predominantly negative. Thus, the product of these two terms is expected to be positive on average. Therefore, it is likely that this contribution will positively contribute to the mean power at each phase.

To obtain the fluctuating component of Eq. 12, Eq. 14 is then phase averaged, where angled brackets are placed only around the terms that have a *ϕ* dependence:

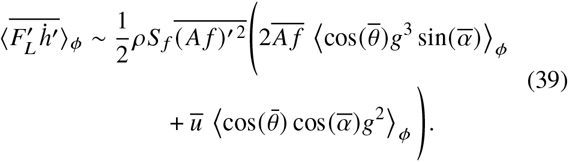

Given the assumptions made previously, over the phase, 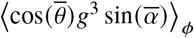 oscillates between approximately 0 and 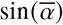. Therefore, its phase average is expected to be an *O*(1) fraction of 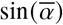, and a value of 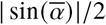 is a reasonable estimate since the oscillation is roughly symmetric. Likewise, 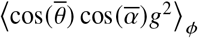 can be approximated as 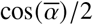.

The mean component of the total power in Eq. 12 can also be approximated as, using the relationship 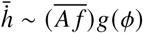:

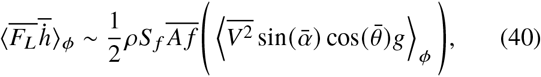

where averages are placed only around the terms that have a *ϕ* dependence. Based on previous assumptions, 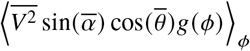 is again expected to be a *O*(1) fraction of 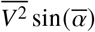, and a value of 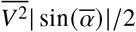 is a reasonable approximation.

Substituting Eq. 39 and 40 into Eq. 12 with the order of magnitude estimates yields Eq. 15.

## Ethics

Animal holding and experimental procedures were approved by the Harvard Animal Care IACUC Committee (protocol number 20-03-3).

## Artificial Intelligence (AI) declaration

AI software was used solely to improve the English language and readability of the manuscript.

## Data Accessibility

Data and all codes utilized to generate the figures contained in this article will be publicly available in Zenodo.

## Authors’ Contributions

Conceptualization: M.A.C, D.R., Y.Z., G.V.L., R.N.; Investigation: D.R., Y.Z., G.V.L.; Formal Analysis: M.A.C., D.R., C.P., Methodology: M.A.C, D.R., Y.Z., C.P., G.V.L, R.N.; Funding Acquisition: G.V.L., R.N.; Project Administration: G.V.L., R.N.; Visualization: M.A.C.; Writing-original draft: M.A.C.; Writing-review&editing: M.A.C., D.R., Y.Z., C.P., G.V.L., R.N.

